# High prevalence of neurocognitive disorders observed among adult people living with HIV/AIDS in Southern Ethiopia: a cross-sectional study

**DOI:** 10.1101/416362

**Authors:** Megbaru Debalkie Animut, Muluken Bekele Sorrie, Yinager Workneh, Manaye yihune Teshale

**Affiliations:** Arbaminch University College of Medicine and Health Sciences Department of Nursing, Arbaminch, Ethiopia; Arbaminch University College of Medicine and Health Sciences Department of public health, Arbaminch, Ethiopia; Bahir Dar University College of Medicine and Health Sciences School of Nursing, Bahir Dar Ethiopia

**Keywords:** People living with HIV/AIDS, HIV-associated neurocognitive disorder, Gamo Gofa, Ethiopia

## Abstract

**Background:** Comprehensive care given to people living with HIV/AIDS is improving from time to time; however, their concurrent cognitive illness is still ignored, under screened and treated particularly in developing countries. And this problem is also striking in Ethiopia. Therefore, the objective of this study was to assess HIV-associated neurocognitive disorders and associated factors among adult people living with HIV/AIDS.

**Methods:** An institution based cross sectional study was conducted in Gamo Gofa zone public Hospitals from April to May, 2017. The systematic random sampling technique was used to select a total of 697 people living with HIV/AIDS. Data was collected using structured interviewer administered questionnaire and International HIV Dementia Scale was used to screen HIV-associated neurocognitive deficits. Data was entered using Epidata version 3.1 and analyzed using SPSS version 20. Both bivariable and multivariable logistic regression analyses were performed to identify associated factors. A P value 0.05 with 95% confidence level was used to declare statistical significance.

**Result:** A total of 684 study participants were included with a 98 % response rate. From the total participants, 56% were females while 44% were males. The mean (±SD) age of the participants was 38.8±8.8years.

The prevalence of HIV-associated neurocognitive disorder was 67.1% (95%CI; 63.6, 70.5). The multivariable logistic analysis indicated that body mass index 16 kg/m2 (AOR 4.149 (1.512-11.387)), being married (AOR 0.9 (0.604-0.623), unemployment (AOR 5.930 (3.013-11.670) and being in WHO clinical stage T3 category (AOR 2.870 (1.098-7.500) were the key predictors of HIV-associated neurocognitive disorders among people living with HIV/AIDS.

**Conclusion:** In this study the prevalence of HIV-associated neurocognitive disorder is higher than the earlier reports in Ethiopia and Africa. The associated factors also vary from that of earlier studies. This indicates the need for formulating preventive mental health programs and policies for people living with HIV/AIDS.

## Background

One of the most devastating health problems facing the human race in the 21 century is the Acquired Immuno-deficiency Syndrome (AIDS) caused by the Human Immunodeficiency Virus (HIV) infection (7). The Sub-Saharan African (SSA) countries have the highest prevalence of people with this disease, accounting for 80% of the cases, despite the continent constituting only 11–12% of the world’s population (2, 7). HIV-associated neurocognitive disorder (HAND) is one of the most common complications of Human Immune Deficiency Virus (HIV) infection (2, 3). It is characterized by cognitive, motor, and behavioral alterations secondary to the preferential impairment of sub cortical structures by the HIV virus (4). The burden of HIV associated neurocognitive disorder (HAND) in adult patients ranges from 19% to 52% in developing countries (9, 10), and from 14% to 64% in low resource settings (8, 11). Even though early detection and follow up of patients believed to be the golden opportunity to increase HIV infected patients’ survival and quality of life, achieving this goal is a major problem in SSA(12). Accessing anti-retroviral therapy (ART) and early linkage of HIV infected individuals in low and middle income countries are a big problem so far. More-over the late presentation of patients in the follow-up clinic makes them vulnerable to HAND and other HIV related complications (9, 10, 13).

HIV-associated neurocognitive disorder causes different problems for HIV infected individuals like poor adherence, alcohol addiction and unable to attend their follow up properly. And this will lead them to develop serious life treating opportunistic infections in the meantime (11, 12). A study done in Singapore, Nigeria, Cameroon, Botswana, Malawi and Dessie Ethiopia indicated that being female, old age and low educational status is significantly related with HIV associated neurocognitive disorder (2, 8, 11-16), whereas, a study done in Brazil, Singapore and Northern Nigeria indicated that CD4 count less than 500 cell/mm3 was associated with HIV-associated neurocognitive disorder(1, 2, 13, 14). Moreover, a study done in South Africa in 2013 showed that highly active anti-retroviral treatment(HAART) naïve and late clinical stage of the illness was factors affecting HIV-associated neurocognitive disorder (10, 16). Body mass index, depression and alcohol abuse were also found to be associated with HIV-associated neurocognitive disorder in Uganda (7). In addition to the above factors, presence of opportunistic infections and poor medication adherence were associated with HIV-associated neurocognitive disorder (17, 18).

In Ethiopia, particularly in the study area, there was no information regarding the prevalence of HIV-associated neurocognitive disorder and its associated factors among peoples living with HIV/AIDS though there are many people living with HIV/AIDS.

Therefore, this study will help to determine the prevalence and the most important associated factors which have an impact on HIV-associated neurocognitive disorders among peoples living with HIV/AIDS.

## Methods

### Study design, setting and area

An institution based cross-sectional study was conducted from April to May 2017 in the Gamo Gofa zone, which is located at a distance of 505 km from Addis Ababa (the capital of Ethiopia). It is administratively organized in15 Woreda, 2 administrative towns, 34 urban and 452 rural kebeles. The total populations are 2,040,972 according to the 2017 Zonal report. There are 3 public hospitals, 73 health centers and 471 health posts in the Gamo Gofa zone currently functional according to unpublished zonal health officials report 2017.

### Study population

All people 18-64 years living with HIV/AIDS (PLHIV) who were getting ART service in Gamo Gofa zone public Hospitals were targets of the study. All selected PLHIV who were getting ART service during the data collection period were included in the study. PLHIV who were seriously ill during the study period were excluded from the study.

### Sample size and sampling procedures

The sample size was determined by taking the two significant variables from previous studies which was CD4 count less than 500 cells/mm^3^ and having a primary educational level or less in a study done at South Wollo, Ethiopia(15, 18) with 80% power, odds ratio 0.36, 95% CI and a none response rate of 10%. Based on this assumption, the final sample size was 697. Systematic random sampling was employed to select study participants. From the total of 3 Hospitals, the samples were allocated to each Hospital proportionally based on their number of PLHIV. Finally, PLHIV in every 3^rd^ PLHIV from their regular follow up in the hospital was enrolled in the study by systematic random sampling method.

### Data collection tools and procedures

Data was collected by face to face interview using an interviewer-administered pre tested and Amharic version (local language of the community) of the International HIV Dementia Scale (IHDS) by 6 trained bachelors of nursing professionals and supervised by the principal investigator. The IHDS is a screening measure of neurocognitive impairment that includes: memory registration for four common objects; motor speed involving the rapid tapping of the thumb and the first digit of the non-dominant hand; speed of information processing and/or executive functioning measured by repetition of a three-position alternating hand sequence; and memory recall of the four objects (11).

This tool has been validated in Sub-Saharan African countries, particularly in South Africa, Cameroon, Botswana, Uganda and Ethiopia and found to have good psychometric properties in African populations with sensitivity of 88% and 80% and specificity of 50% and 55% respectively at a cutoff 10 or less (7, 8, 10, 11). So, in this study cutoff point <=9.5 was used. It is brief, inexpensive and can be administered easily by other health workers, not necessarily by a physician (2).

#### Adherence of patients to their medication was assessed using Morsiky-8 item scale (MMAS)

The MMAS-8 is a self-report questionnaire with 8 questions (items). Items 1 through 7 have response choices “yes” or “no” whereas item 8 has a 5-point Likert response choices. Each “no” response is rated as “1” and each “yes” is rated as “0” except for item 5,in which each response “yes” is rated as “1” and each “no” is rated as “0”. For item 8, if a patient chooses a response “0”, the score is “1” and if they choose response “4”,the score is “0”. Responses “1, 2, 3” are respectively rated as “0.25, 0.75, 0.75”. Total MMAS-8 scores can range from 0 to 8 and have been categorized into three levels of adherence: high adherence (score=8), medium adherence (score of 6 to < 8), and low adherence (score< 6) (18, 23).

#### Alcohol Use Disorder Identification Test

The Alcohol Use Disorder Identification Test (AUDIT) is a 10-item instrument designed by the World Health Organization to screen excessive consumption, hazardous, and harmful use of alcohol. The instrument has been validated in many countries, including Sub-Saharan African countries. A score of 8 and above is indicative of either harmful use or dependence(2). And a cut of point >=8 was used in this study.

#### Drug Abuse Screening Test

Drug Abuse Screening Test (DAST) instrument is designed to screen for drug use other than alcohol(24).The DAST-10 version was used in this study and is answered in a “yes or no’’ pattern. A score of 3 and above is indicative of a problem with psychoactive substance abuse(2).

Other clinical parameters which were important for the objective was extracted from patients chart.

### Data quality management

Data collectors were trained for two days by doing standardization exercise in order to minimize errors.

The questionnaire was developed in English and then translated into Amharic (local language) and back to English then review was made for consistency of translation of the language. Data were collected after pretest has been conducted on 35 (5%) of PLHA from non-selected health institutions. The principal investigator made day to day on-site supervision during the whole period of data collection. The collected data were reviewed and checked for completeness, accuracy and consistency by investigator.

### Data management and analysis

Data entry and cleaning was made using Epidata 3.1 and exported to SPSS software package version 20 for analysis.

Cross-checking and data cleaning was carried out by running frequencies of each variable. For specific objective one, descriptive statistical methods such as frequencies, percentages, proportion with 95% C.I has been used. Mean and standard deviation was also used to summarize various characteristics of the participants.

For specific objective two, cross tabulation and bivariable logistic regression were used to explore the relation between the outcome variable and the different independent variables using crude odds ratio with 95% C.I. Finally, to determine the independent factors associated with the outcome variable, multivariable logistic regression model was done, and presented using adjusted odds ratio (AOR) with 95% confidence interval. Variables with P-value < 0.2 of the bivariable analysis were taken in to the multivariable logistic regression model. Model fitting was checked using log likelihood and Hosmer-Lemeshow test. Finally, variables with P < 0.05 in the multivariable analysis were considered as significant.

## Results

### Socio-demographic characteristics

A total of 684 people living with HIV/AIDS were participated in this study, making the response rate of 98%. The mean age of the respondents was 38.8±8.8 years. Among the study participants majority 383 (56.0%) were females and 283 (41.4%) were widowed (Table 1).

**Table 1.** Distribution of PLHA by their socio-demographic characteristics at Arbaminch general hospital, Chencha & Saula district hospitals ART clinic, 2018.

| Variables |  | Frequency (n=684) | Percent |
| --- | --- | --- | --- |
| Sex | Male | 301 | 44.0 |
|  | Female | 383 | 56.0 |
| Age | 18-25 | 27 | 3.9 |
|  | 26-35 | 224 | 32.7 |
|  | 36-45 | 255 | 37.3 |
|  | 46-56 | 164 | 24.0 |
|  | >56 | 14 | 2.0 |
| Marital status | Single | 143 | 20.9 |
|  | Married | 142 | 20.8 |
|  | Widowed | 115 | 16.8 |
|  | Divorced | 284 | 41.5 |
| Residence | Urban | 595 | 87.0 |
|  | Rural | 89 | 13.0 |
| Religion | Orthodox | 370 | 54.1 |
|  | Protestant | 247 | 36.1 |
|  | Muslim | 61 | 8.9 |
|  | Others | 6 | 0.9 |
| Educational status | Unable to read & write | 130 | 19.0 |
|  | Able to read & write | 206 | 30.1 |
|  | Primary | 217 | 31.7 |
|  | Secondary & above | 131 | 19.2 |
| Occupation | Employed | 655 | 95.8 |
|  | unemployed | 29 | 4.2 |
| Monthly income(ETB) | ≤585 | 166 | 24.3 |
|  | 586-1650 | 263 | 38.5 |
|  | 1651-3145 | 181 | 26.5 |
|  | ≥3146 | 74 | 10.8 |

Key: ETB=Ethiopian Birr (Ethiopian currency).

### Clinical status

Most of the participants 588 (86.0%) were in WHO clinical stage T1 category and their mean CD4 count was 610 ± 278 cells/mm^3^. And 95.6% of the participant had no any history of central nervous system related opportunistic infections. The majority of the participants 643 (94.0%) were low adherent to their ART treatment (Table 2).

**Table 2.**
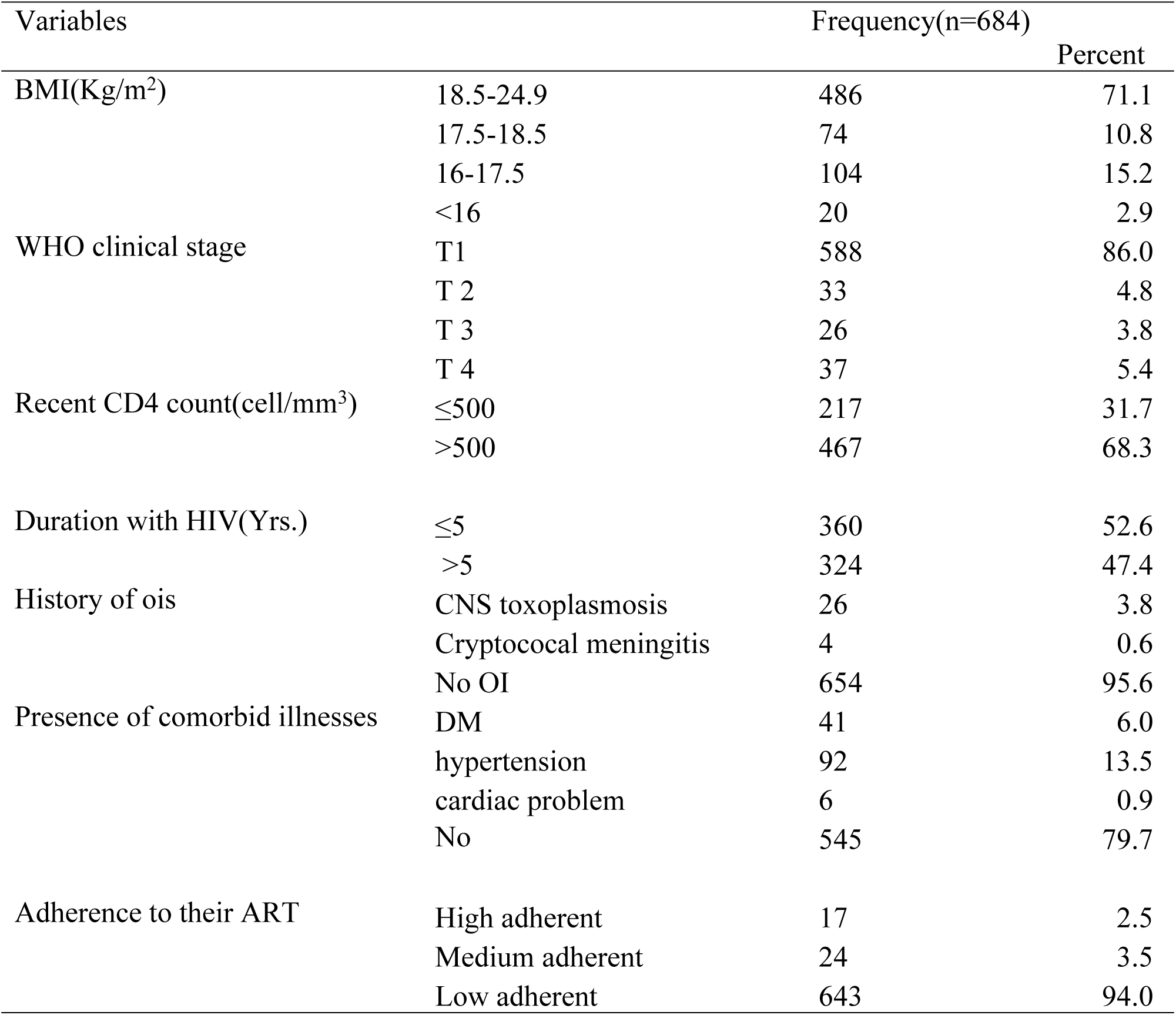
Distribution of PLHA by their clinical status at Arbaminch general hospital, Chencha & Saula district hospitals ART clinic, 2018.

### Alcohol and other drug usage pattern

Alcohol use disorder identification test was used to screen participants in this tool and the score is meant to be classified into two cut off points as less than eight (never abuse alcohol) and greater than or equal to eight (which screens alcohol abuse) and in this study majority 462 (67.6%) of the participants were never abuse alcohol. Drug abused screening test (DAST-10) was also used to screen for drugs other than alcohol usage and most of the participants 678 (99.1%) of them were scored between 0-3 which is non-drug abuse.

### Prevalence of HIV associated neurocognitive disorder (HAND)

Among the study participants who have been screened for HAND, 459 (67.1% (95%CI; 63.6, 70.5) screened as having HAND. Procedural implementation of international HIV dementia scale (IHDS) was as follows: The first measurement on IHDS was timed finger tapping, on this part motor speed was assessed, 65 (9.5%) were scored 4 out of 4 which is normal. On the second part psychomotor speed measurement was assessed, of whom 109 (15.9%) had performed the sequential procedure, scoring 4 out of 4. Finally, memory recall was assessed and was found at 174 (25.4%) had recalled all the four items without any clue scoring 4 out of 4.

### Factors associated with HAND

Bivariable and multivariable analysis showed that the association between HIV-associated neurocognitive disorder and marital status, body mass index, WHO clinical status and occupational status were significant. After controlling /adjusting other variables like (sex, alcohol abuse, religion, educational status, CD4 count, drug abuse, duration with the virus, medication adherence, monthly income, comorbid illness, opportunistic infection history). The odds of HAND among married participant decrease by 10% compared to those with unmarried/single participants (AOR=0.9 CI (0.604-0.623)).

In this study the likelihood of having the risk of HAND among PLHA was four times in those participants who had a body mass index of less than 16 kg/m^2^ than those participants who had a body mass index of 18.5-24.9 kg/m^2^ (AOR 4.149 CI (1.512-11.387). The odds of having HAND among unemployed participants were almost six times higher than those who were employed (AOR=5.930 CI (3.013-11.670)). Similarly, being in WHO clinical staging T3 category were approximately three times more likely to have the risk of HAND (AOR =2.870 CI (1.098-7.500)) when compared to those with WHO clinical staging T1 category (table 3 below).

**Table 3.** Bivariable and multivariable analysis showing the association between socio-demographic and clinical related variables with HAND among PLHA in Arbaminch general hospital, Chencha & Saula district hospitals ART clinic, 2018.

| Variables |  | HAND |  | COR | AOR |
| --- | --- | --- | --- | --- | --- |
|  |  | YES | NO |  |  |
| Marital status | Single | 58 | 85 | 1 | 1 |
|  | Married | 28 | 114 | 0.578(0.454-0.736) | 0.9(0.604-0.623)* |
|  | Divorced | 104 | 180 | 0.246(0.162-0.371) | 0.449(0.234-0.860) |
|  | Widowed | 35 | 80 | 0.438(0.294-0.651) | 0.671(0.346-1.298) |
| Body mass index | 18.5-24.9 | 162 | 324 | 1 | 1 |
|  | 17.5-18.5 | 17 | 57 | 0.298(0.174-0.513) | 0.598 (0.319-1.120) |
|  | 16-17.5 | 33 | 71 | 0.465(0.308-0.702) | 1.270(0.749-2.154) |
|  | <16 | 13 | 7 | 1.857(0.741-4.655) | 4.149(1.512-11.387)* |
| Occupational status | Employed | 183 | 441 | 1 | 1 |
|  | Unemployed | 42 | 18 | 2.333(1.343-4.053) | 5.930(3.013-11.670)* |
| WHO clinical stage | T1 | 193 | 395 | 1 | 1 |
|  | T2 | 11 | 22 | 0.500(0.242-1.031) | 1.291(0.541-3.081) |
|  | T3 | 12 | 14 | 0.857(0.396-1.853) | 2.870(1.098-7.500)* |
|  | T4 | 9 | 28 | 0.321(0.152-0.681) | 1.039(0.429-2.517) |

Key: \* = $p < 0.05$ , Number of total participants n=684.

## Discussions

Determination of the prevalence and associated factors of HIV-associated neurocognitive disorders among peoples living with HIV/AIDS by institutional based cross sectional study in southern Ethiopia public hospitals was the main focus of this study. At the base of this, the prevalence of HAND among people living with HIV/ AIDS was 67.1%. And IHDS demonstration was highly associated with marital status, body mass index, occupational status and WHO clinical stage category.

The finding of this study was in line with the study done in Sub Saharan African population which showed the prevalence of HAND (64.4%) (7), and in Kenya (65.6%) (25).

However, the prevalence of HAND in the current study was higher than a cross-sectional study done in Brazil (53%)(16), (53%)(4), in Botswana (38%)(8) Indonesia (51%)(26), Nigeria 30%(20), South Africa (25%) (27), South Asia (22.7%) (13) and in Ethiopia at Debre Markos North West (24.8%) (17), Dessie (36.4%) (18) and the Mekele referral hospital Northern Ethiopia 33.3% (21). The differences might be accounted to the neurovirulence strain differences and sociodemographic differences. In addition, most of the above studies used smaller sample size than this paper. Moreover, educational status can also play a role in the performance of IHDS since; in this study unable to write was 19%, which might increase the prevalence of HIV associated neurocognitive disorder. Above all the difference in the care of patients with HIV in different settings might be different and will contribute to the difference in prevalence.

On the other hand, this study was lower than study done in Bamenda, Cameroon (85%), a cross-sectional study at Bamenda Regional Hospital AIDS-treatment Centre among 297 PLHA who have been on antiretroviral therapy (11).

The discrepancy in the prevalence rate might be due to the neurovirulence clade differences for the more common clade in other sub Saharan countries is A&D but in Ethiopia is not clearly known merely clade C thought to be the prevalent one which is neurotropic viruses like A&D but less likely to be associated with a neurocognitive disorder than clade A&D. The differences in the predictors for the development of HAND might also have an effect for the discrepancy since in the previous study the common predictor was stated as having primary education or less and having HIV symptoms but not in the current study. And the discrepancy can also be explained by sample size difference, in previous study lesser sample size, which is 297 much less than the current study. In addition, other possible reasons for the differences between the present and other studies were mainly due to differences in the method used, differences in study populations, using different cutoff points in different studies, some inappreciable cultural and educational difference, and environmental factors. Above all discrepancies might also be explained by the application of the IHDS which might need well-educated participant.

This study found out that, marital status was significantly associated with the development of HAND; married participants were less likely to develop HAND than single individuals. This may be due to married couples may motivate each other to exercise, eat healthfully, maintain social ties and smoke and drink less. All these things could prevent the development of HAND. In single, widowed or divorced participants’ stress level will be high which further can affect nerve signaling in the brain and impair cognitive abilities. In addition, people who are married tend to be financially better and stable than others, a factor that is closely associated with many aspects of human health.

In this study, body mass index of <16 kg/m^2^was significantly associated with HAND which was nearly in line with USA Baltimore/Los Angeles sites of the Multicenter AIDS Cohort Study (28). This is due to, body mass index less than 16 kg/m^2^ or sever adult malnutrition can increase risk of neurocognitive disorder. Perhaps not surprisingly, the risk of different kind of neurocognitive disorder/dementia can be increased by vitamin deficiencies, specifically B-12, and other forms of Vitamin B which seems to be the vitamin most likely to potentiate the development of dementia if not consumed in adequate quantities. Malnutrition in PLHA causes severe immune suppression so that they can be at risk of developing many central nervous system opportunistic infections in cortical or subcortical structure of the brain which facilitates neurocognitive disorder or dementia.

In the current study occupational status had significant association with HAND. Being unemployed were at risk of developing HAND than employed one. This study is nearly similar to a cross sectional study done among 361 participants in Asia(29), a cohort study done in Thai(30) and cross sectional study among 604 participants in Ireland and Irish ART Clinic (31). Employment needs any person to acquire new skills, participate in different organization and engage socially, all of these can give meaning to a person’s life and provide income. All of these activities can boost a person’s cognitive health. Unfortunately, if they get unemployed, they will lose or limit at least employment driven cognitive health benefits. Especially in people with HIV/AIDS money is everything for them without it, they will be exposed to different opportunistic infection due to lack of money to buy and take different prophylactic drugs. In the present study, there was statistically significant association between WHO clinical staging and HAND. This finding is supported by other a study done among 268 patients, demonstrate that this factor, was the predictive variables in the development of HAND (32). During late stage of the disease, the virus will be very high and can cause different central nervous system related opportunistic infection. In addition the virus itself can replicate and multiply in different parts of the brain. Both cases will facilitate development of HIV associated neurocognitive disorder.

Generally; the results of this study provide additional information on HAND particularly in the local areas where this study is done. The result of this study can also give some information on specific variables which affect HAND, science these variables were not mentioned well in other previous studies. Above all, the finding of this study will contribute about the burden and risk factor of HAND to the scientific world.

### Limitation of the study

Despite including participants from three hospitals it might not be applicable for all PLHA in Ethiopia. This study included only patients on highly active antiretroviral therapy so being on HARRT might underestimate the prevalence of neurocognitive disorder.

### Conclusion

There is a high prevalence of HIV-associated neurocognitive disorder than that reported earlier in Ethiopia and Africa. The associated factors also vary from that of earlier studies. Therefore, this study indicates the need for formulating preventive mental health programs and policies for patients with HIV/AIDS.

**List of Abbreviations**
AOR: Adjusted Odds Ratio
AIDS: Acquired Immune Deficiency Viruses
ART: Anti-retroviral therapy
BMI: Body Mass Index
CDC: Center for Disease Prevention and Control
CI: Confidence
COR: crude odd ratio
HAART: Highly Active Antiretroviral Therapy
HAND: HIV-Associated neurocognitive disorder
HIV: Human Immuno deficiency virus
IHDS: International HIV Dementia Scale
PLHA: People Living With HIV/AIDS
SSA: Sub Saharan Africa
WHO: World Health Organization.

## Ethics approval and consent to participate

Ethical clearance was obtained from an Ethical Review Board of College of Medicine and Health Sciences, Arba Minch University & permission letter was obtained from Gamo Gofa Zonal health department and respective hospitals.

The nature of the study was fully explained to the study participants to obtain oral consent in the study and any information was kept confidential.

Based on their informed consent, participants had full right to refuse or discontinue in the study without any compromise in the services they get from each facility. For that purpose, one-page consent paper attached as a cover page to each questionnaire. Any personal identifiers were not included in the questionnaire. Any participant, who was screened as having HIV-Associated neurocognitive disorder, was recommended to be evaluated further for possible treatment and follow up evaluations back to the clinicians.

## Consent for publication

Not applicable

## Availability of data and materials

Data of this study will be obtained by contacting the corresponding author via this or Tel no. +251920049549

## Authors’ contributions

MDA: conceived the study, coordinated the overall activity, carried out the statistical analysis and drafted the manuscript.

MBS: participated in the design of the study, reviewing the drafted manuscript.

YW: participated in the design of the study, reviewing the drafted manuscript.

MYT: participated in the design of the study, reviewing the drafted manuscript.

All authors read and approved the final manuscript

## Acknowledgements

The authors would like to thank Arbaminch University. Furthermore, we extend our heartfelt gratitude to Gamo Gofa zone Health Bureau; we also want to thank all respondents, data collectors for their active participation during the data collection process. The last but not the least we acknowledge Mr Getaneh Alemu for his constructive comment and language editing.

